# Identification and Characterisation of Proteins Binding to a G-Quadruplex Origin G-rich Repeated Element in Mammalian Cells

**DOI:** 10.1101/2023.03.30.534946

**Authors:** James R. A. Hutchins, Isabelle Peiffer, Serge Urbach, Jean-Louis Mergny, Philippe Marin, Domenico Maiorano, Marcel Méchali

## Abstract

In metazoan cells, replication of genomic DNA initiates from thousands of discrete chromosomal loci known as origins. Proteins such as the Origin Recognition Complex (ORCs) associate with origins, but this does not show clear sequence specificity for DNA binding. Genome-wide origin mapping studies have shown that the region surrounding the replication initiation site contains motifs such as the Origin G-rich Repeated Element (OGRE), proximal to the majority of origins. Here, using an approach coupling DNA affinity purification to quantitative proteomics, we identified proteins that interact specifically with an OGRE. Three of the top-scoring interactors, Dhx36, Pura and Tial1, were selected for further study. We show that Dhx36 and Tial1 localise to the nucleus and their knockdown decreased cells in S-phase resulting in their accumulation in the G_1_ phase of the cell cycle. Altogether these results indicate that these OGRE-binding factors may play roles in DNA synthesis in mammalian cells.

## 1. Introduction

In eukaryotes, replication of genomic DNA is initiated at thousands of discrete chromosomal loci, known as replication origins. Genome-wide mapping studies in metazoan cells have identified thousands of origins distributed throughout the chromosomes (Fragkos *et al*, 2015; Prioleau & MacAlpine, 2016). These studies showed that although metazoan origins do not share a strict DNA sequence consensus motif, they are preferentially associated with a number of chromosomal characteristics, which comprise a combination of genetic and epigenetic factors (Prioleau & MacAlpine, 2016; Hutchins *et al*, 2016). Origins are significantly associated with GC-rich DNA, CpG islands and transcriptional start sites (Cadoret *et al*, 2008; Sequeira-Mendes *et al*, 2009; Cayrou *et al*, 2011). Epigenetic elements observed at replication origins were mainly the presence of the histone variant H2A.Z, Polycomb marks, and a nucleosome-free region at the G-rich element (Cayrou *et al*, 2015). The application of sophisticated machine-learning algorithms to sets of human origins from developmental and pathological states has enabled the prediction of a common set of “core” origins active in different cell types (Akerman *et al*, 2020).

DNA replication origins are bound by the origin recognition complex (ORC) that in turn recruits Cdc6 and Cdt1 proteins, thus allowing recruitment of the hexameric Minichromosome Maintenance (MCM2-7) protein complex, to form pre-replicative complexes (pre-RCs), which then progress to Initiation Complexes (in G_1_ phase), until their activation at the entry to S-phase (Hu & Stillman, 2023). Replication origins are in excess in the genomes of eukaryotes. Not all origins bound by pre-RC proteins do eventually fire; indeed, only a part of them are activated at each cell cycle, no more than 30% in metazoan cells. This flexibility the choice of origins to be activated is a main feature of eukaryotic replication origins (Méchali, 2010).

One distinctive motif associated with the majority of mammalian replication origins is the Origin G-rich Repeated Element (OGRE), identified first in the mouse and *Drosophila* genomes (Cayrou *et al*, 2011; Cayrou *et al*, 2012). In mice, a majority of OGREs was found to contain the consensus motif G_(3)_-N_(1-7)_-G_(3)_-N_(1-7)_-G_(3)_-N_(1-7)_-G_(3)_ (where N is any base), compatible with the formation of G-quadruplex (G4) structures (Cayrou *et al*, 2012). These non-Watson-Crick DNA formations arise when a set of four guanine bases (forming a planar structure called a G-tetrad) stack to form a stable structure (Bochman *et al*, 2012). G4s are enriched at certain chromosomal sites, including telomeres and promoter regions of genes (Rhodes & Lipps, 2015). This observation of motifs containing potential G4s in the vicinity of replication origins was subsequently extended to human cells (Guilbaud *et al*, 2022; Besnard *et al*, 2012). Experiments in which G4-forming sequences upstream of replication origins were either ablated or displaced to other loci provided evidence that they do contribute to the functionality of replication origins (Prorok *et al*, 2019; Valton *et al*, 2014). Two main features of the OGRE motif are its location upstream of the replication initiation site and its nucleosome-free nature (Cayrou *et al*, 2012; Cayrou *et al*, 2015). These characteristics may suggest a regulatory function as a “landing pad” for proteins that establish pre-RCs in the vicinity of the initiation site.

With the aim of identifying OGRE-associating proteins as potential candidates for origin recognition in metazoans, we performed a proteomic screen in which proteins from a nuclear extract were isolated using an immobilised G4-containing model OGRE sequence, followed by their identification by quantitative mass spectrometry. Three proteins showing specificity for the OGRE/G4 were selected for functional studies, Dhx36, Pura and Tial1. Their characterisation provides evidence for their implication in DNA replication.

## 2. Results

### Affinity purification of OGRE/G4 interactors

To identify proteins associating with a G4-containing OGRE, we adopted a proteomic approach, combining DNA affinity purification (DAP) with quantitative mass spectrometry (MS). For this DAP-MS method, we chose a “model” OGRE derived from the mouse genome, from which the highest quality genome-wide origin mapping data were available (Cayrou *et al*, 2011; Cayrou *et al*, 2015) and which was supported by the strongest evidence that it forms a G4. Our desired model OGRE required the following criteria:

(1) it should be a G-rich region proximal to a verified origin of replication, at a distance from the initiation site (IS) consistent with the genome-mean distance between OGREs and ISs; (2) it should have a sequence fitting the consensus motif for G4s, and to have a high propensity score for G4 formation using a leading prediction algorithm; (3) there should be evidence that this DNA sequence forms a G4 *in vitro*.

For the first criterion, a list of OGREs were derived from replication origin mapping data from mouse chromosome 11, generated via the nascent-strand (NS) mapping method (Cayrou *et al*, 2011). For the second criterion, we narrowed the list of OGREs to those whose sequence fits the classical consensus for forming a G4 structure: G_(3)_-N_(1-7)_-G_(3)_-N_(1-7)_-G_(3)_-N_(1-7)_-G_(3)_. The likelihood of these OGREs to form G4s was calculated using the G4Hunter algorithm (Bedrat *et al*, 2016), which scores DNA sequences for G4 propensity on a scale from -4 (least likely) to +4 (strongest likelihood). A score of +2 or greater indicates a very strong propensity to form a stable G4. A shortlist of OGREs with score +2 or greater was derived. For the third criterion, biophysical properties of high G4Hunter-scoring OGREs were tested based on two complementary assays of OGRE-containing oligonucleotides. The circular dichroism (CD) spectrum of a DNA molecule shows a characteristic pattern of peaks and troughs at certain molecular ellipticity values when a G4 is present (Del Villar-Guerra *et al*, 2018). A Thermal Difference Spectrum (TDS), obtained by measuring ultraviolet spectra of a DNA molecule above and below its melting temperature, shows characteristic features that indicate the presence of a G4 (Mergny *et al*, 2005).

Taking into consideration these three criteria, the best-scoring candidate OGRE was “OGRE3”, which is located close to the 3′ end of the first intron of the *Rai1* (retinoic acid induced 1) gene on mouse chromosome 11, and 274 bp proximal to the nearest origin (**Figure 1**). OGRE3 has the G-rich sequence GGGGGCGGGGAGGGAAGGGGG, which fits well with the canonical G4 consensus motif, and has a high G4Hunter score of +3.05. CD and TDS spectra were previously reported for this sequence (Prorok *et al*, 2019), which was known as “Ori1” in that study. Genomic editing experiments with this OGRE sequence showed that its presence and position can contribute to replication activity within a genomic locus, indicating its functionality.

**Figure 1.**
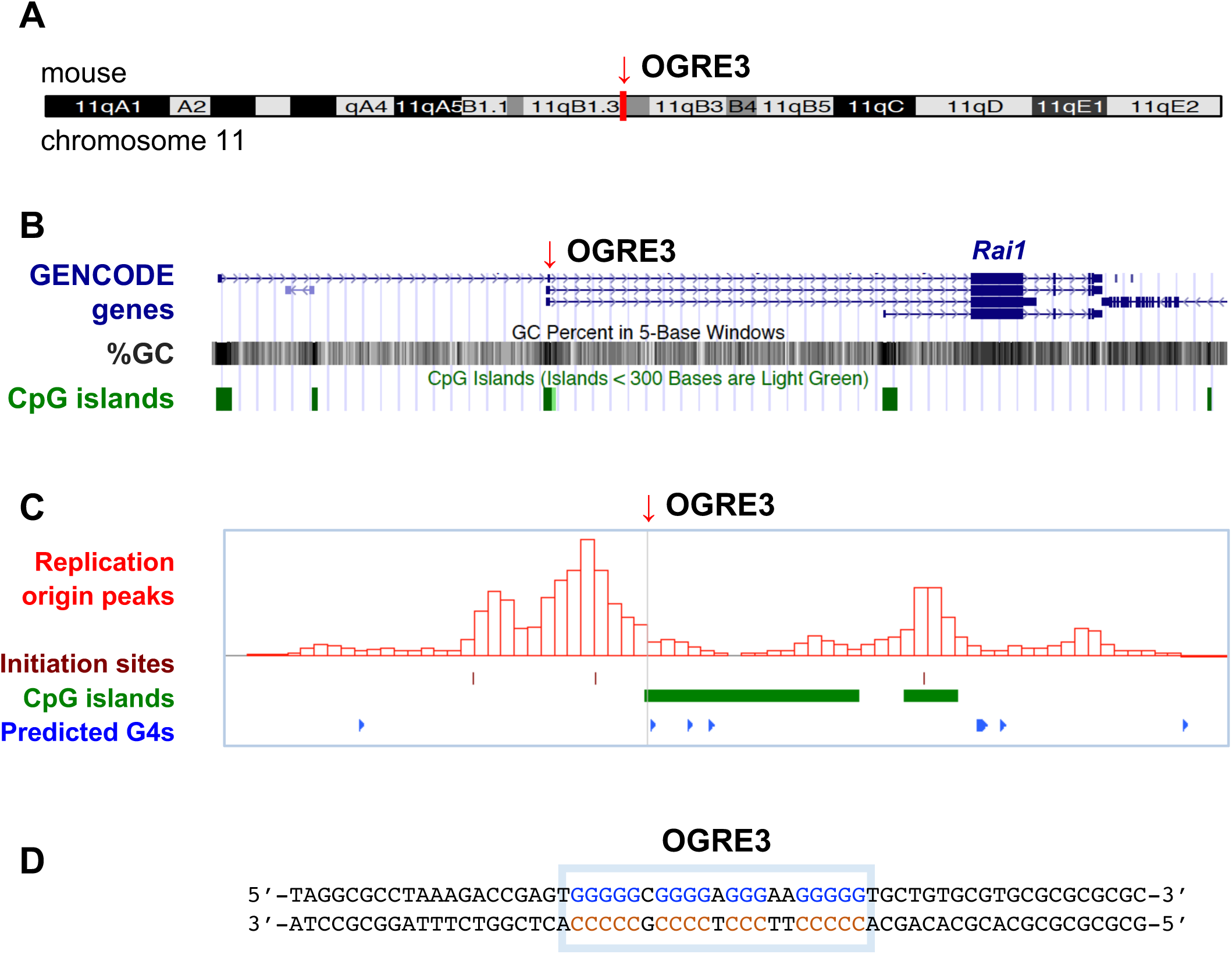
– Characteristics of OGRE used for DAP-MS. **(A)** Location of OGRE3 within the mouse genome. **(B)** Characteristics of the genomic environment local to OGRE3. **(C)** Position of OGRE3 relative to mapped replication origins, CpG islands and predicted G4s. **(D)** Sequence of OGRE3 (within blue box).

In order to isolate and characterise proteins associated with OGRE3, we adopted and adapted the DNA affinity purification mass spectrometry procedure described by Nordhoff and colleagues (Nordhoff *et al*, 1999) that we refer to as DAP-MS (**Figure 2**). Here, proteins from a cell extract are captured by immobilised DNA molecules of a specific sequence, then analysed by mass spectrometry for identification and relative quantification. As OGRE3 is of mouse origin, we chose NIH 3T3 cells (immortalised mouse fibroblasts) as the source material for proteins.

**Figure 2.**
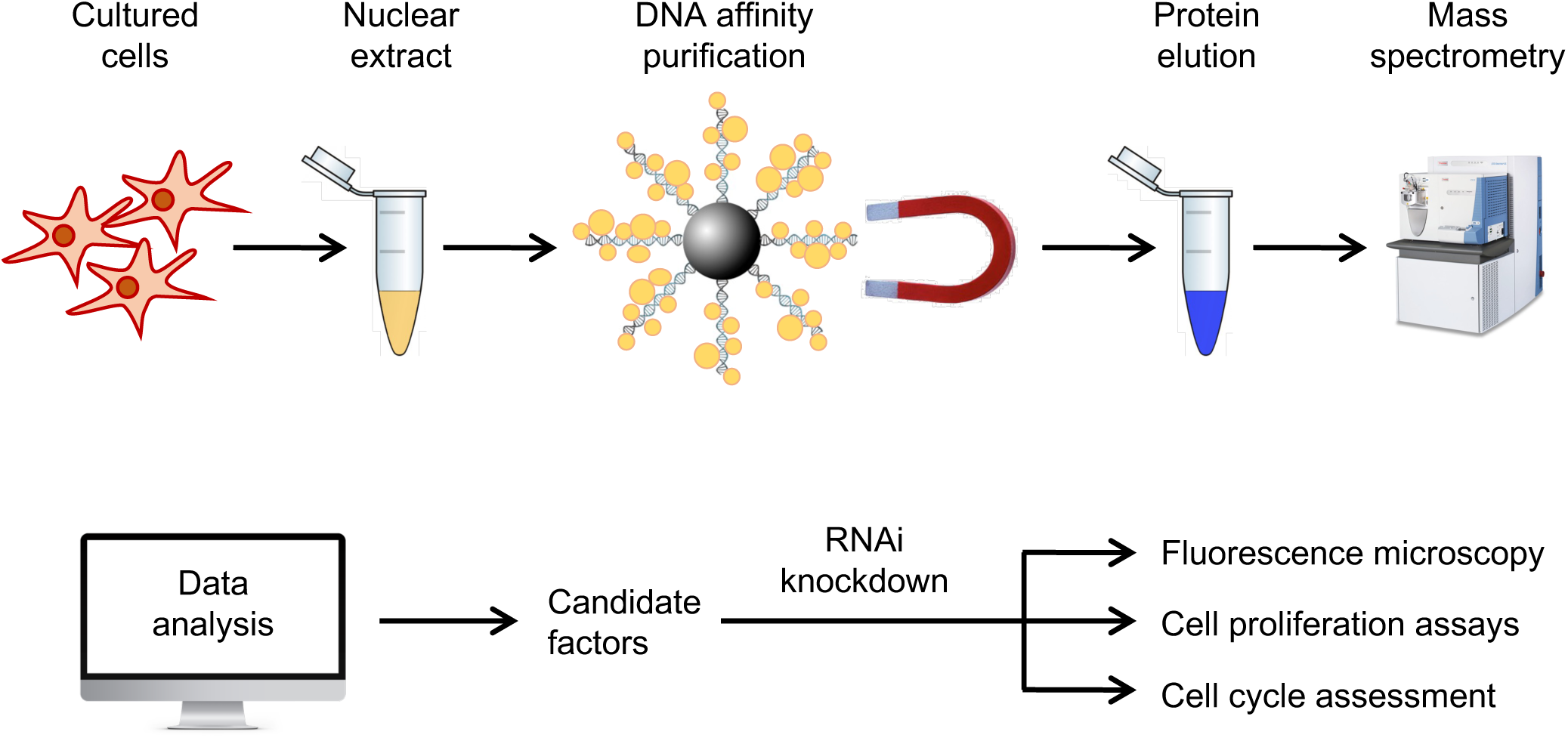
– Workflow for identification and characterisation of proteins showing preferential binding to OGRE/G4s. Nuclear extracts were prepared from cultured mouse cells, and proteins isolated using DNA Affinity Purification (DAP) using a DNA probe containing a well-characterised OGRE. Label-free mass spectrometry and data analysis were used to identify and quantify proteins, and to single out leading candidate factors for functional studies. Expression of candidate factors in cells was knocked down using RNAi, and resultant phenotypes characterised using fluorescence microscopy, cell proliferation assays and flow cytometry-based cell cycle assessments.

### Isolation of OGRE-binding factors by DAP-MS

In a first set of DAP-MS experiments, we designed a DNA probe of mouse genomic sequence containing the OGRE3 sequence at its centre. In terms of length of flanking sequences, we wished to give sufficient “landing pad” for a sizeable protein complex. For this, our starting point was by analogy with transcription complexes, which may span the typical 25-30 bases from the TATA box to the transcription start site (Kornberg, 2007). Based on this, our DAP probe was 80 bp, with the OGRE3 sequence (21 bp) flanked on both sides by 29 bp and 30 bp of genomic sequence.

In addition to the wild-type sequence (OGRE3^WT^), we designed a mutated form (OGRE3^MUT^) that maintains the G/C base composition of the OGRE, but which lacks contiguous stretches of guanines, ensuring that it cannot form a G4 (**Figure 3A**).

**Figure 3.**
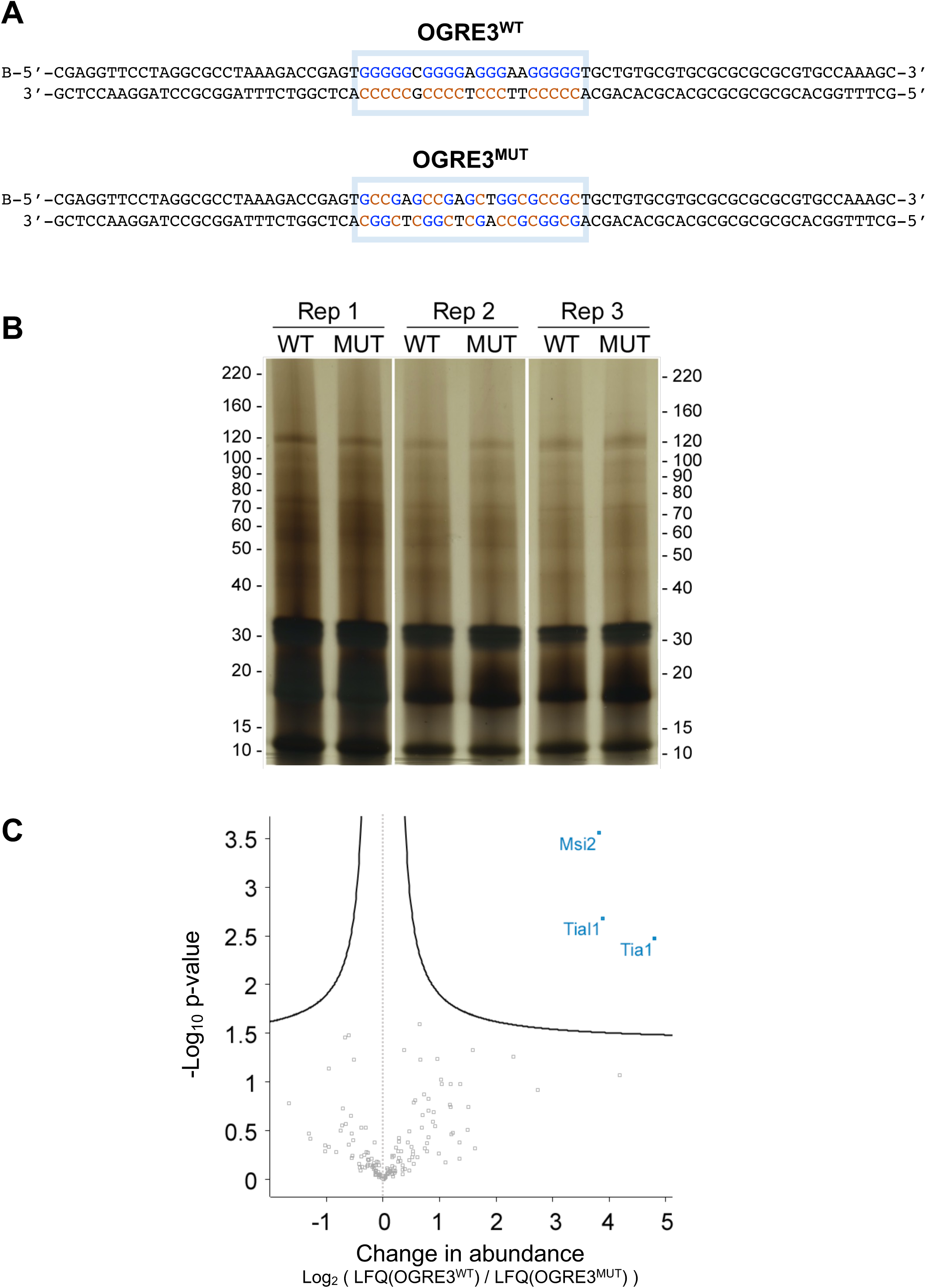
– Identification of proteins bound to 80-bp OGRE3. **(A)** Sequence of 80 base-pair OGRE3^WT^ and OGRE3^MUT^ -containing DNA probes. B indicates a biotin moiety. OGRE portions are highlighted by blue boxes. **(B)** Proteins from nuclear extracts of NIH 3T3 cells were isolated on beads coupled to OGRE3^WT^ or OGRE3^MUT^, as indicated, in three replicates (Rep 1, Rep 2, Rep 3). A sample of each purification was separated by SDS-PAGE then the gel was silver stained; the positions of protein molecular mass markers (kDa) are shown. **(C)** Volcano plot indicates proteins identified by DAP-MS as showing specific preference for the 80-bp OGRE3^WT^ DNA probe. The *x*-axis shows log_2_ change in abundance of proteins with OGRE3^WT^ compared to OGRE3^MUT^; proteins located to the right of the dotted line are enriched on OGRE3^WT^. The *y*-axis shows -log_10_ of the p-value; proteins to the upper right of the solid curved line surpass the confidence threshold, and are significant.

DNA affinity purification was performed using this probe in three separate replicates, following which 10% of the proteins isolated was analysed by SDS-PAGE and silver staining to assess the yield and quality of the preparation (**Figure 3B**). The remainder of the preparation was analysed by LC-MS/MS, then mass spectral data were analysed to identify protein hits matching to the mouse UniProt database (UniProt Consortium, 2023). Entries corresponding to proteins from well-established lists of non-specific binding proteins were filtered from the list.

Determination of LFQ intensities allowed the median fold differences in the abundances of proteins identified with the OGRE3^WT^ and OGRE3^MUT^ conditions to be calculated. Further statistical analysis revealed which proteins surpassed the false discovery rate (FDR) threshold, i.e. were statistically significant hits, as shown in a volcano plot (**Figure 3C**).

From this experiment, three proteins surpassed the significance threshold: Tia1, Tial1, and Msi2. Tia1 (T-cell-restricted Intracellular Antigen-1), and Tial1 (Tia-Like 1, also known as nucleolysin Tiar) are highly similar proteins. Tial1 modulates gene expression by affecting RNA transcription, splicing, stability and translation (Reyes *et al*, 2009). Both also play functional roles in RNA quality control in stress granules (Anderson & Kedersha, 2008). In terms of nucleic acid binding features, both Tia1 and Tial1 contain three RNA Recognition Motif (RRM) domains, with sequence-specific binding to DNA also documented (Suswam *et al*, 2005). Tial1 has been linked to cell cycle control, as it is essential for the G_2_/M checkpoint, accumulating in nuclear foci in late G_2_ and prophase in cells undergoing replication stress (Lafarga *et al*, 2019). Msi2 (Musashi homologue 2) is a transcriptional regulator protein, containing two RRM domains, that targets genes involved in development and cell cycle regulation (Fox *et al*, 2015). Elevated levels of this protein are associated with poor prognosis in certain cancer types (Kharas & Lengner, 2017).

In a second DAP-MS experiment, we considered the possibility that the 80-bp length of the DAP-MS probe may have limited the accessibility of proteins to the OGRE3 sequence. Thus we designed a longer, 120-bp DAP-MS probe, this time with 50 bp of genomic DNA flanking the OGRE3 sequence, again in wild-type (OGRE3^WT^) and mutated (OGRE3^MUT^) forms (**Figure 4A**). DAP-MS was performed using this probe from nuclear extracts of NIH-3T3 cells as described previously, with 10% being analysed by SDS-PAGE and silver staining (**Figure 4B**). In this case, the specificity of the remaining proteins for OGRE3^WT^ compared to OGRE3^MUT^ was determined based on two metrics: the difference between LFQ intensities for proteins associating with OGRE3^WT^ and OGRE3^MUT^, and the ratio between these two values. Thirty-seven proteins yielded a positive difference, i.e. showed an increased abundance on OGRE3^WT^ compared to OGRE3^MUT^, indicating that they had shown specificity in binding OGRE3^WT^ (**Figure 4C** and **Supp Table 1**). In terms of LFQ intensity ratios, 31 proteins had a value greater than one, however only three of these (Pura, Purb, Tial1) had a ratio greater than two. Six proteins (Dhx36, Fam114a1, Lims1, Arfgap3, Rrp12, Lactb) were identified in the OGRE3^WT^ but not in the OGRE3^MUT^ condition (**Figure 4C**).

**Figure 4.**
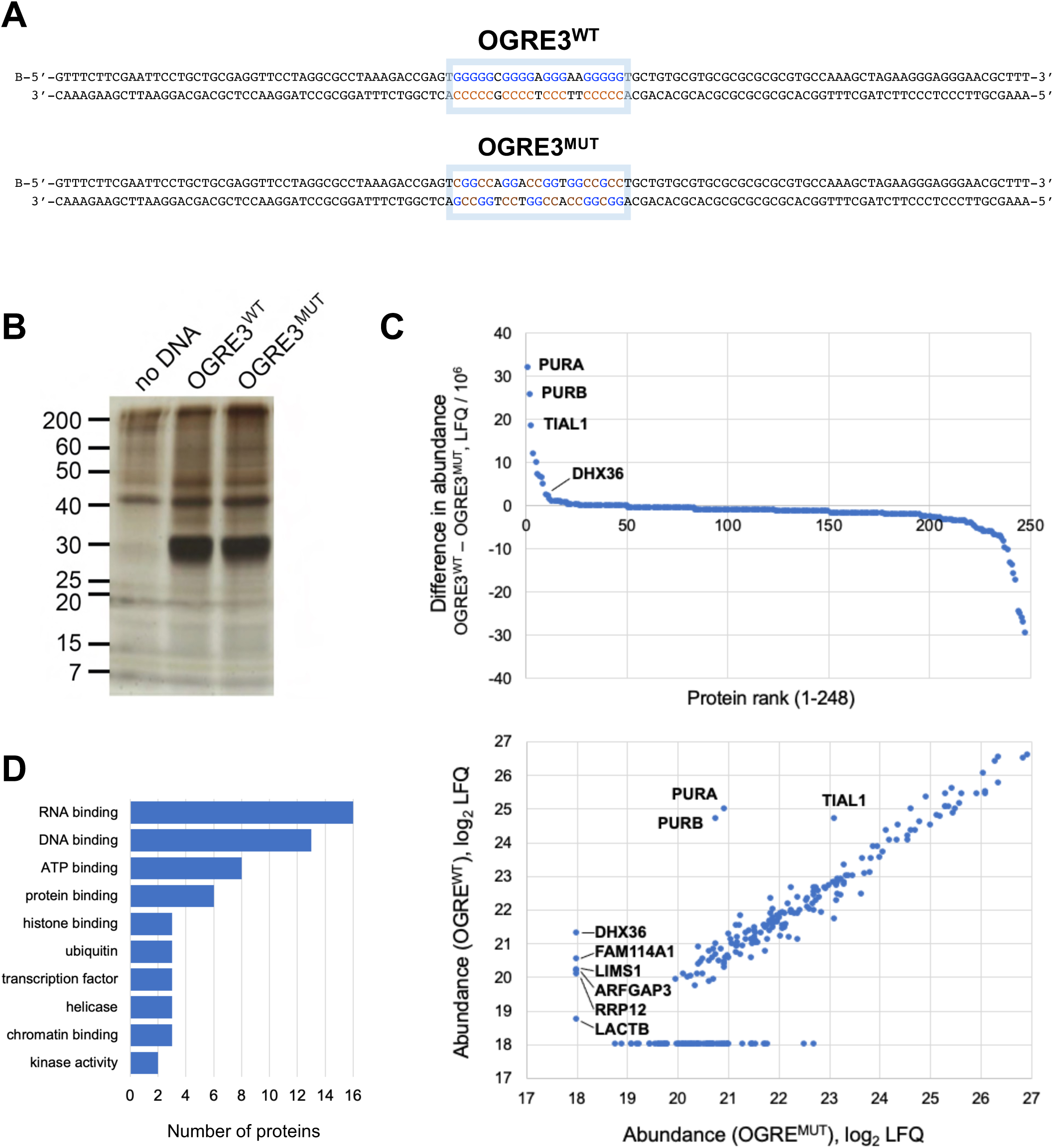
– Identification of proteins bound to 120-bp OGRE3. **(A)** Sequence of 120 base-pair OGRE3^WT^ and OGRE3^MUT^ -containing DNA probes. B indicates a biotin moiety. OGRE portions are highlighted by blue boxes. **(B)** Proteins from nuclear extracts of NIH 3T3 cells were isolated on beads coupled to OGRE3^WT^ or OGRE3^MUT^, or no DNA, as indicated. A representative purification is shown. A sample of each purification was separated by SDS-PAGE then the gel was silver stained; the positions of protein molecular mass markers (kDa) are shown. **(C)** Above: plot of difference in abundance between OGRE3^WT^ and OGRE3^MUT^, for the 248 proteins identified by DAP-MS (after filtering out non-specific proteins). Below: plot of log abundances of OGRE3^WT^ versus OGRE3^MUT^. Proteins not identified under a condition were arbitrarily assigned a value of 18. **(D)** Gene Ontology Molecular Function terms corresponding to the 37 proteins showing preference for OGRE3^WT^ and OGRE3^MUT^. The ten most popular terms are shown.

### Investigating properties of the proteins identified

In order to determine which of the 37 proteins identified by DAP-MS as showing specific binding to OGRE3^WT^ may have the most potential interest for follow-up studies, previously determined properties and functions of these factors were investigated by systematic querying of publicly-accessible databases (Hutchins, 2014).

The Gene Ontology (GO) database was queried to ascertain whether the proteins identified show nuclear localisation or have established roles in the cell cycle or DNA replication. Of the 37 proteins, 28 have GO Cell Compartment annotations indicating evidence for localisation to the nucleus (including the nucleolus) or chromatin. Of those, 20 proteins have Molecular Function annotations indicating evidence for nucleic acid (DNA or RNA) binding activity (**Supp Table 1, Figure 4D**). GO Biological Process terms showed that two of the 37 proteins, DNA ligase 3 (LIG3) and TAR DNA-binding protein 43 (TARDBP) had entries related to cell cycle control, and three proteins Pur-alpha (Pura), Lig3 and Histone-binding protein RBBP7 (Rbbp7), have reported roles relating to DNA replication.

A further source of genes shown to have cell cycle roles is from a genome-wide siRNA-based knockdown screen that identified 1249 human factors required for cell division (Neumann *et al*, 2010). Only one of the 37 proteins, heterochromatin protein 1-binding protein 3 (HP1BP3) had been identified as a factor required for cell division in this screen.

One may surmise that if a factor plays an important role in such a fundamental process as DNA replication or cell division cycle control then it would be essential, i.e. gene deletion of the gene or ablation of the protein would result in cell death. In the case of ORC, all subunits are indeed essential, although incomplete replication may lead to cell cycle arrest in mitosis rather than S-phase (Chou *et al*, 2021). To assess how essential each of the 37 factors are, effects of their knockout in mice, where recorded, were retrieved from the Mouse Genome Informatics (MGI) database (Blake *et al*, 2021). 12 of the 37 proteins are encoded by genes whose knockout leads to embryonic lethality, 2 lead to postnatal lethality, and 15 lead to postnatal abnormalities or pathologies (**Supp Table 1**).

### Choice of OGRE interacting factors for functional experiments

We wished to select a small number of high-confidence candidate factors from the list of 37 for functional experiments. Our choice was made based on the criteria of specificity for G4, as judged by differential abundance in our DAP-MS screens; the presence of features consistent with activity in the nucleus and with nucleic acid binding, and possible clues to involvement in G4 binding and/or DNA replication from the literature. The three factors we chose were Dhx36, Pura and Tial1, for the reasons given below.

Dhx36 (DEAH-Box Helicase 36) was the highest-scoring protein identified in the OGRE3^WT^ but not in the OGRE3^MUT^ condition, underlying the specificity of its interaction with the G4-containing probe. Dhx36 is an ATP-dependent helicase with G4 resolving activity (Vaughn *et al*, 2005), and a crystal structure of Dhx36 in a complex with G4 DNA reveals the mode of action of its binding and unwinding (Chen *et al*, 2018). The presence of Dhx36 in the list of OGRE interactors served to validate our differential DAP-MS approach, as well as providing an additional high-confidence G4-binding candidate for functional studies.

The next two proteins showing the greatest LFQ intensity differences were Pura and Purb. Pura was originally described as the “Pur Factor” a protein binding a purine-rich element (*PUR*) proximal to mammalian origins of replication (Bergemann & Johnson, 1992). Pura, Purb and Purg constitute a paralogous trio in human cells, each containing a tandem array of three “PUR” domains, conserved from bacteria to humans, capable of binding DNA or RNA. Pura is reported to be implicated in regulating the transcription of several genes, in the response to replicational stress (Wang *et al*, 2007), and to play a role in a number of neurological conditions (Daniel & Johnson, 2018).

After Pura and Purb, the protein with the highest LFQ intensity differential was Tial1. This protein can bind DNA and RNA, thus could interact with an OGRE either directly, or indirectly via other proteins or nucleic acids. Tial1 was the only protein identified as strongly positive in both DAP-MS screens, and was thus chosen for functional studies.

#### Downregulation of OGRE-binding factors slows down cell proliferation

To assess whether the three candidate factors have roles in controlling the rate of cell proliferation, and whether they affect cell cycle progression, their expression was knocked down separately in asynchronous U2OS cells using a two-step transfection procedure: cells, in triplicate, were transfected with siRNAs against the candidates (or a luciferase negative control, siLuc), and three days later transfected again with the same. Cell proliferation was assessed at days 3 and 6, and at day 6 the cells were pulse-labelled with the nucleotide analogue BrdU, allowing ongoing DNA synthesis to be assessed by flow cytometry. Western blotting of cell proteins at days 3 and 6 post-transfection showed that expression of Dhx36 was completely repressed, and that of Pura and Tial1 greatly reduced (**Figure 5A**).

**Figure 5.**
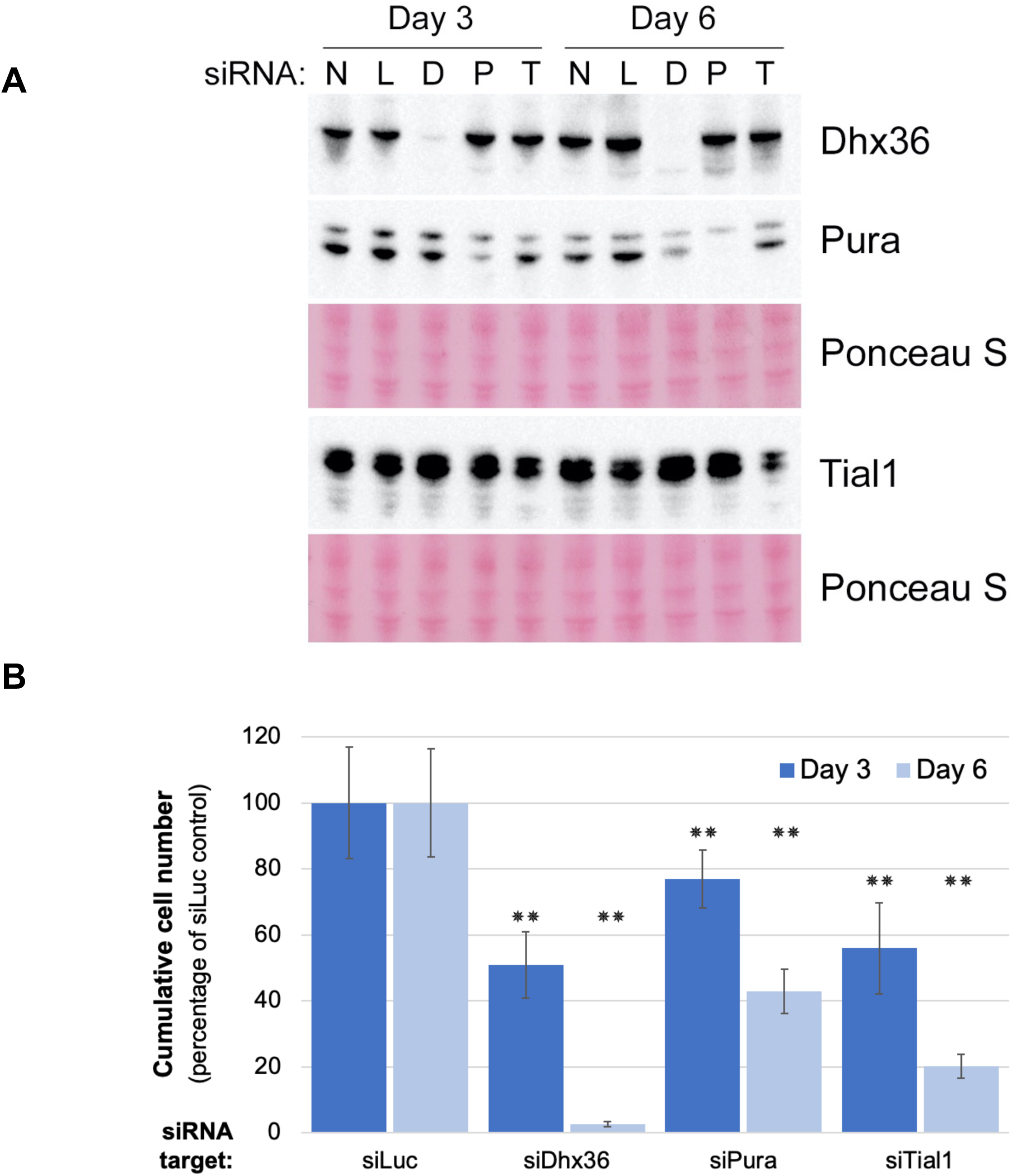
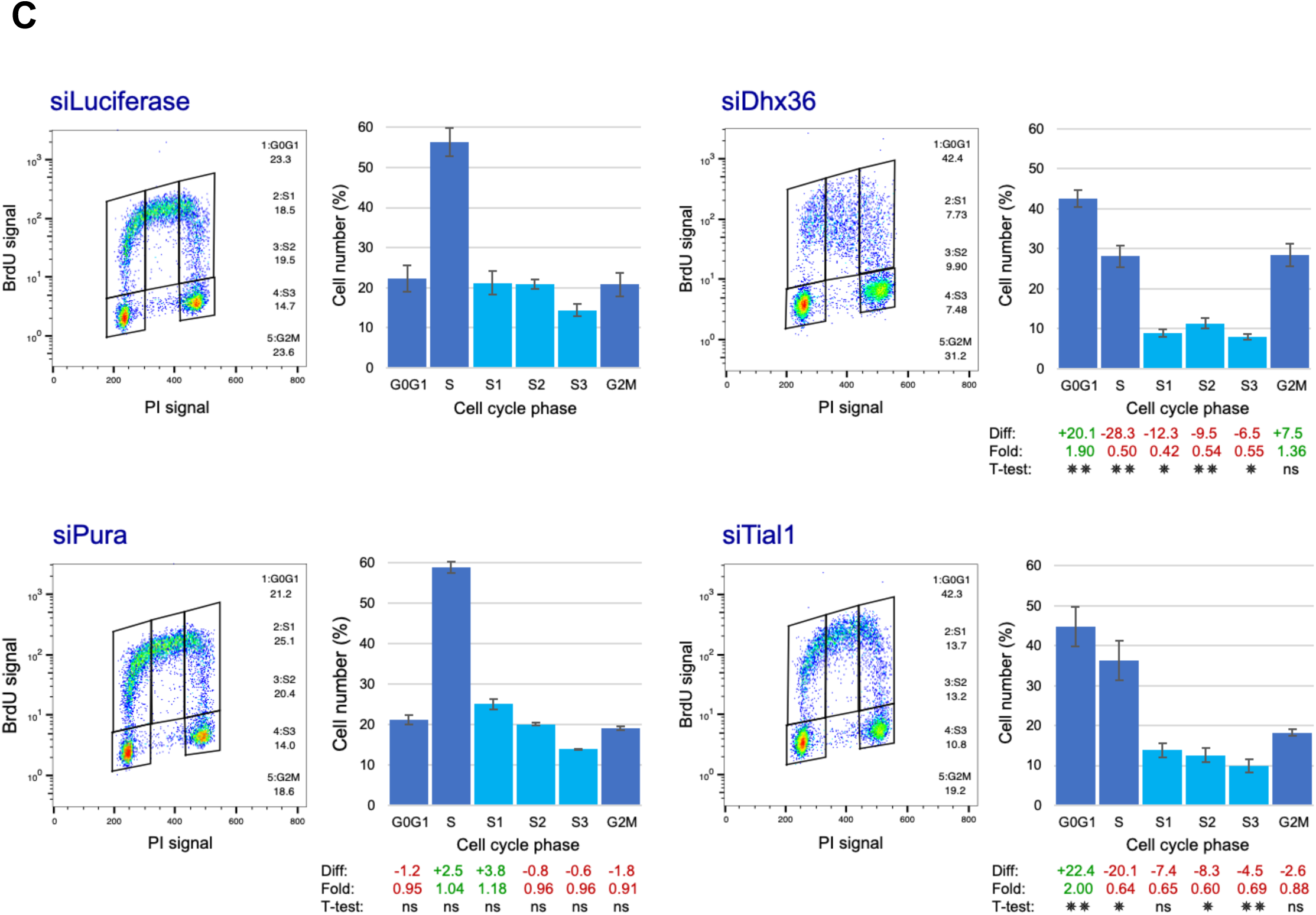
- Knockdown of candidate factors affects cell proliferation and cell cycle progression. A two-stage RNAi protocol was employed, in which triplicate plates of U2OS cells were transfected with siRNAs targeting the factors. Non-transfected cells and siLuciferase-transfected cells served as negative controls. Three days later, cells were counted, re-seeded and re-transfected with siRNAs as before. At day 6, cells received a pulse of BrdU, were counted, and samples taken for western blotting and fixed for analysis by flow cytometry. **(A)** Western blot showing knockdown of the indicated factors. RIPA extracts were prepared from cells harvested at days 3 and 6, as shown, 10 µg protein were loaded per lane, and the membrane immunoblotted with antibodies to the proteins indicated. N, non-transfected cells; L, siLuciferase; D, siDhx36; P, siPura; T, siTial1. Ponceau S-stained membrane portions show equal loading of protein on the gels. **(B)** Cumulative cell numbers at day 3 and day 6 post-transfection with siRNAs targeting candidate factors are shown as percentages of the cell count for the control siLuciferase (siLuc) transfections. Error bars represent standard deviations. Statistical significances were determined using Student’s t-test; **, p ≤ 0.005. **(C)** Two-dimensional flow cytometry plots of PI (DNA content) and BrdU incorporation (DNA replication) of cells at day 6 post-transfection. Gates indicate delimitations of cell sub-populations in defined cell cycle phases. S-phase was divided into S_1_ (early-S), S_2_ (mid-S) and S_3_ (late-S) sub-phases. Quantifications of the phases are shown in column plots. The differences (Diff) and fold changes (Fold) in the mean cell population for each phase in a given siRNA condition compared to that for siLuciferase are shown (green, increase; red, decrease). Error bars represent standard deviations. Statistical significances were determined using Student’s t-test; ns, not significant; *, p ≤ 0.05; **, p ≤ 0.005.

Downregulation of each of the three factors resulted in a significant reduction in cell proliferation. In particular, the number of cells in the siDhx36 condition was reduced to 51% (compared to the siLuc control condition) after 3 days and less than 3% after 6 days. For siPura, the cell count was reduced to 77% after 3 days and 43% after 6 days. For Tial1, the number was reduced to 56% after 3 days and 20% after 6 days (**Figure 5B**). These results indicate that all three of these OGRE/G4-binding factors are required for cells to proliferate at the normal rate.

Assessment of cell cycle phase distribution of cells at day 6 was performed by two-dimensional flow cytometry. Here, the preparation protocol enabled a high degree of separation between BrdU-negative and BrdU-positive cells, and enabled S-phase to be subdivided into three sub-phases: early-S, middle-S and late-S, referred to as S_1_, S_2_ and S_3_, respectively (**Figure 5C**). For the control (siLuc) condition, 21% of cells were in G_1_ or G_0_ phases (G_0_/G_1_), and 20% were in G_2_ or M phases (G_2_/M), whereas the proportion in S-phase was about 55%. A breakdown of the S-phase population into S_1_, S_2_ and S_3_ showed a distribution of 20%, 20% and 15%, respectively. In the Dhx36 knockdown condition, the proportion of cells in G_0_/G_1_ phases had doubled to 42%, whereas the percentage in S phase had decreased by half, and that for G_2_/M was increased by about a third, to 28%. Inspection of sub-phases S_1_, S_2_ and S_3_ showed a slightly greater proportion in mid-S than for early and late S-phases. The number of cells in G_2_/M phases was increased by a third. As well as a reduced proportion of cells in S-phase, the level of BrdU signal was reduced as the cells progressed through S-phase, forming a diffuse cloud rather than the distinctive “horseshoe” shape in the 2D flow cytometry plot (**Figure 5C**).

In the Tial1 knockdown condition, cells showed a doubling in G_0_/G_1_ phase relative to siLuc, and a reduction by a third in the number of S-phase cells (with an increased S_1_ and a reduced S_3_). A small decrease in G_2_/M cells was not significant. In the siPura condition, the proportion of cells in the three main cell cycle phases was not significantly changed relative to siLuc, however inspection of sub-phases S_1_, S_2_ and S_3_ showed that the proportion in early S-phase was increased, whereas that for late S was decreased.

Collectively, these experiments showed that out of the three candidate factors studied, all three are necessary for correct proliferation, however only Dhx36 and Tial1 had a significant effect on cell cycle progression, the cells in S-phase being significantly reduced, suggesting a potential role in DNA replication.

#### Subcellular localisation of the factors

As these candidate factors were isolated as binding to the genomic OGRE element, they would be expected to localise in the nucleus of the cell. To verify that this is the case, we knocked down their expression using RNA interference, and assessed their localisation by fluorescence microscopy. Imaging of the cells stained for Dhx36 and Tial1by immunofluorescence showed that both Dhx36 and Tial1 did indeed show predominantly nuclear localisation (**Figure 6**). Quantification of signal intensities showed the nuclear/cytoplasmic ratios of both factors to be around 2. Treatment with siRNA resulted in a reduction in signal intensity in the nuclear compartment for both factors of between 57% and 79%. Concomitantly, the intensities in the cytoplasmic compartment were reduced by 83% to 91% for Dhx36 and Tial1, respectively. We could not assess properly the subcellular localisation of Pura since the antibody did not show a specific reactivity by immunofluorescence, as assessed by comparing staining before and after knockdown by siRNA.

**Figure 6.**
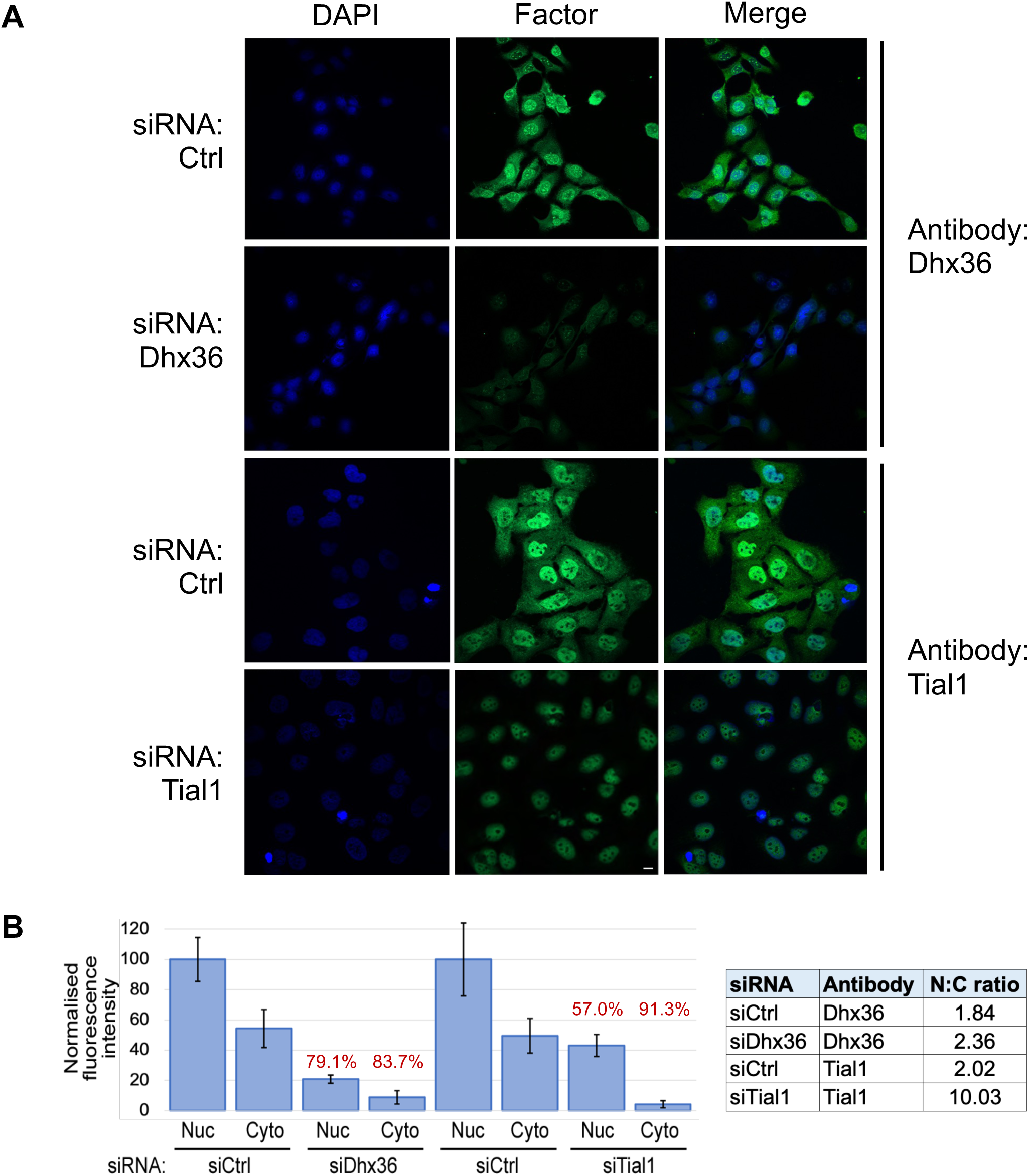
– Subcellular localisation of Dhx36 and Tial1. **(A)** Immunofluorescence microscopy images of U2OS cells transfected with siRNAs against the factors indicated (or a non-specific control sequence, Ctrl), fixed and stained with antibodies against the factors and with DAPI. Scale bar, 10 µm. **(B)** Plot of green intensity values in the nucleus (Nuc) or cytoplasm (Cyto) from (A). Values represent mean measurements from ten cells from each image shown, normalised to the value for the siCtrl transfection in each set. Error bars show standard deviations. Numbers in red refer to percentage reduction relative to siCtrl. The table to the right shows nuclear/cytoplasmic ratio (N:C ratio) values for each condition.

## 3. Discussion

Genomic DNA sequences containing the OGRE motif appear to contribute to origin functionality of replication origins (Prorok *et al*, 2019; Valton *et al*, 2014), however the underlying molecular mechanism is currently poorly understood. In this perspective, we have applied a DNA affinity purification and quantitative proteomic approach to find proteins that specifically bind to OGRE/G4 elements. Maybe surprisingly, we did not identify any known DNA replication factors associated with the OGRE probe. One possible explanation for this could be the observation that OGREs are located not exactly at replication initiation sites, but oriented in nucleosome-free regions upstream of them. Hence, OGREs may have a stimulatory activity on DNA replication origins, through recruitment of specific factors that may modulate DNA and/or chromatin structure, such as those isolated in the screen described in this report. This is reminiscent of the ancillary activity of DNA elements flanking the autonomously replicating sequence (ARS) in the yeast *S. cerevisiae*, such as the A, B and C motifs that bind transcription factors (Dhar *et al*, 2012). Notwithstanding, one cannot exclude the possibility that the OGRE probe we used to pull down binding factors was too short to allow recruitment of known replication factors. Among the proteins identified, we selected three factors, Dhx36, Pura and Tial1, all of which contain nucleic acid binding motifs, that could be potential candidates as proteins important to modulate DNA replication origin activity through OGRE binding.

### Dhx36

The identification of Dhx36 as the highest-scoring protein interacting with OGRE3^WT^ for which no peptide was identified in the OGRE3^MUT^ condition validates our approach to the specific identification of G4 interactors, and opens an avenue of investigation for the possible role of a known G4 helicase in the recognition of DNA replication origins.

Knockdown of Dhx36 slowed down cell proliferation and resulted in a reduction by half of the number of S-phase cells, as well as a reduction in the BrdU signal. This latter result indicates reduced DNA synthesis was taking place, a phenomenon that could be due to a lower rate of origin firing, slower elongation, or a series of replication fork stalling and restarting events. G4s may represent obstacles to the passage of replicative DNA polymerases along the chromatin (Sato *et al*, 2021), and our findings in S-phase are consistent with a possible role of Dhx36 in the initiation of DNA replication, perhaps by contributing to DNA unwinding, although an additional possible role during the elongation step cannot be excluded. The implication of Dhx36 in DNA synthesis is novel, since Dhx36 was originally identified as RHAU, an <u>R</u>NA <u>h</u>elicase isolated in association with the <u>AU</u>-rich elements of mRNAs (Tran *et al*, 2004). The protein, also given the name G4 Resolvase 1 (G4R1), was shown to be the major source of cellular G4 DNA resolving activity of DNA and RNA G4s (Creacy *et al*, 2008; Vaughn *et al*, 2005). Single-molecule studies showed the activity of Dhx36 to be dependent on its nucleotide-bound state (You *et al*, 2017). Pathway analysis using the REACTOME resource (Gillespie *et al*, 2022) also shows a role for Dhx36 in a pathway that acts as a cytosolic sensor of pathogen-associated DNA as part of the innate immune system (**Supp Figure 1A**). KEGG Pathways (Kanehisa & Goto, 2000) analysis indicated a role for Dhx36 in RNA degradation (**Supp Figure 1B**).

Analysis of Dhx36 interactors with the STRING database (Szklarczyk *et al*, 2021) reveals only four partner proteins, including the transcription factor IRF7, the RNA binding protein LIN28A and the DNA helicase Dhx9 (**Supp Figure 2A**). Interestingly, Dhx9 has been previously reported to interact with the subunits of the MCM2-7 protein complex in U2OS cells (Cheng *et al*, 2017), revealing a possible functional partnership in the initiation of DNA synthesis. In conclusion, Dhx36 may have a double role, in DNA synthesis and RNA metabolism.

### Pura

Pura was the highest scoring protein by differential LFQ intensity in the DAP-MS screen. Pura is a member of the Pur gene family, which is highly conserved in eukaryotes. Humans have three paralogous genes, *PURA*, *PURB* and *PURG*, the latter being expressed as two proteins (Purg-A and Purg-B), all four proteins containing three tandem “PUR” domains that show DNA/RNA binding capacity. Pura is a multifunctional protein with diverse functions regulating different aspects of nucleic acid metabolism, including gene transcription, RNA transport and mRNA translation (Daniel & Johnson, 2018; Johnson, 2003). Consistent with having roles regulating gene expression, Pura mutations are present in some neurodevelopmental disorders, including a condition known as PURA Syndrome (Molitor *et al*, 2021). Purb was also identified in our screen, with a similar (but slightly lower) intensity to Pura; Pura and Purb are known to heterodimerise (Kelm *et al*, 1999), thus co-purification may have occurred in this case.

Pura was first reported as a protein binding a *PUR* element, a G-rich sequence proximal to a zone of DNA replication initiation near the *MYC* gene (Bergemann & Johnson, 1992). A recent ChIP-seq study identified *PUR* elements on a genome scale (Shi *et al*, 2021); one motif corresponding to a subset of these sequences bears close resemblance to that of the OGRE (**Figure 7**). This *PUR* motif contains the spaced GGGs required for G4 formation, so one may surmise that a subset of *PUR* elements may form G4s. It has been proposed that Pura may bind G4 DNA (Daniel & Johnson, 2018), although this awaits experimental determination. Pura can unwind both single- and double-stranded DNA (Wortman *et al*, 2005), and crystal structure studies showed modes of binding to tri- and hexanucleotide repeats and the mechanism of its unwinding activity (Weber *et al*, 2016), however if this unwinding applies to G4-containing DNA elements is still unclear.

**Figure 7.**
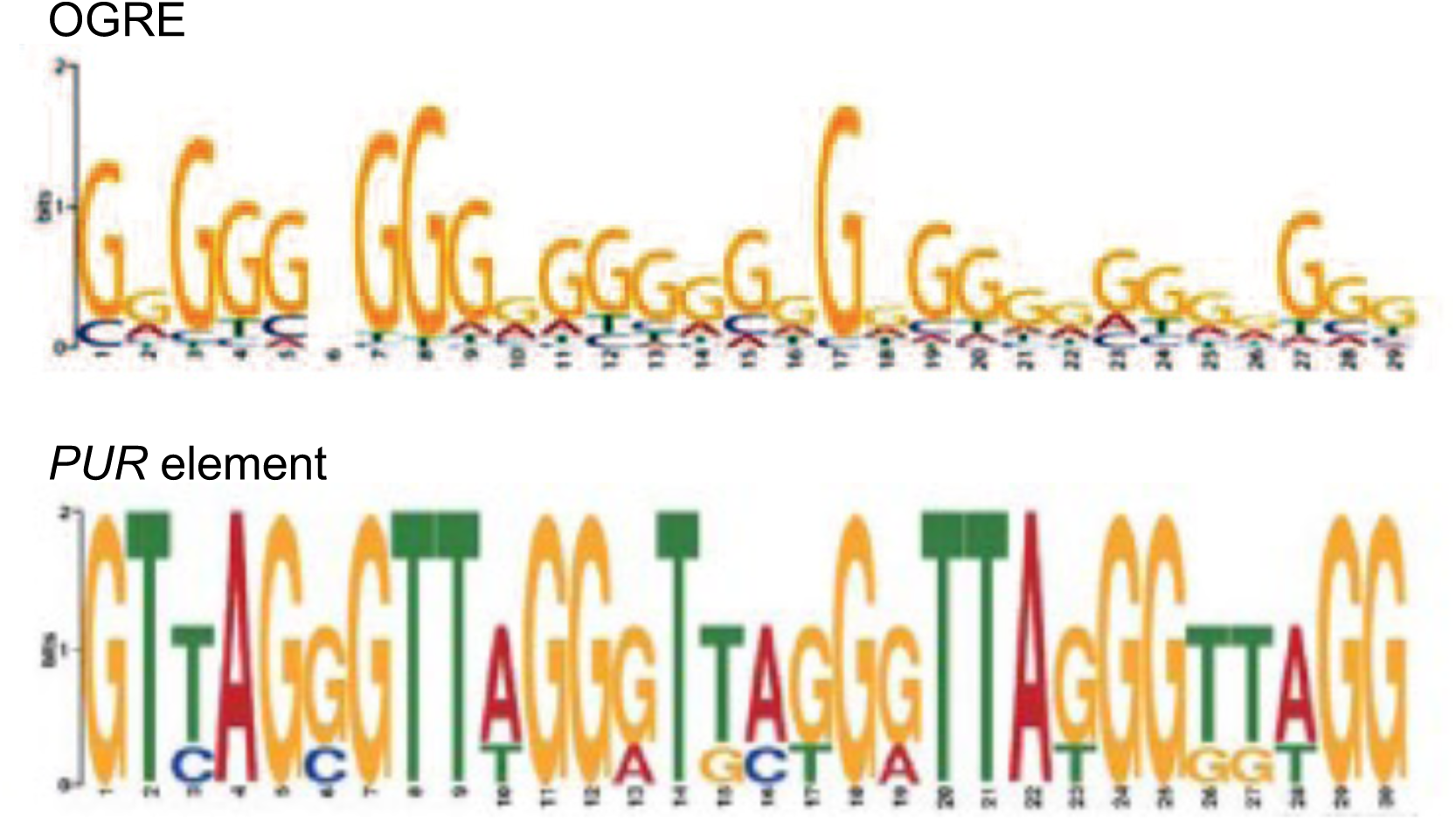
- Comparison of sequence logos from the mouse genome for (above) OGRE and (below) G-rich *PUR* element. OGRE: (Cayrou *et al*, 2012). *PUR* element: (Shi et al, 2021).

In terms of possible roles in regulating DNA replication, STRING analysis shows Pura to interact with ten proteins, including nine factors involved in DNA replication: DNA polymerases alpha-1, alpha-2 and delta-3, PCNA, RPA subunits 1,2,3 and 4, and the replisome-associated protein DONSON (Zhang *et al*, 2020) (**Supp Figure 2B**). Pura has been linked with cell cycle control in that it binds several regulators of cell proliferation, including the retinoblastoma protein RB1, E2F-1, Sp1, YB-1, cyclin T1/Cdk9, and cyclin A/Cdk2 (White *et al*, 2009).

Taken together, these observations suggest that Pura could have a role analogous to a DNA helicase in binding to sequence elements in the genome (including OGREs). However, we could not observe a significant effect of PURA downregulation on the cell cycle or S-phase progression. This result can be explained by redundancy between Pur family members and/or with other DNA helicases, including Dhx36, at OGREs.

### Tial1

Tial1, the third-highest scoring hit in our OGRE/G4 interaction screen after Pura and Purb, was originally discovered as a cDNA with a similar sequence to TIA (<u>T</u>-cell-restricted <u>I</u>ntracellular <u>A</u>ntigen-1 cytotoxic granule associated RNA binding protein) (Kawakami *et al*, 1992), and called TIAR for TIA-related protein.

Tial1 is a member of a family of RNA-binding proteins, has three RNA recognition motifs, and binds adenine and uridine-rich elements in mRNA and pre-mRNAs of a wide range of genes. It regulates various activities including translational control, splicing and apoptosis. Tial1 is well conserved in eukaryotes, including invertebrate metazoans and plants (https://www.ncbi.nlm.nih.gov/homologene/?term=TIAL1). It is an essential gene in mouse, the knockout showing an embryonic lethal phenotype (Beck *et al*, 1998).

Consistent with Tial1 having multiple roles in RNA metabolism, analysis of interactors with the STRING database reveals nine Tial1 partners including two heterogeneous nuclear ribonucleoproteins, two nuclear cap-binding proteins and RNA-binding protein 25 (**Supp Figure 3**). Pathway analysis using REACTOME shows a role for Tial1 in mRNA splicing: together with Tia1 and other proteins it contributes to the formation of the FGFR2 IIIb splice variant that is characteristic of epithelial cells (**Supp Figure 3**).

Our knockdown of Tial1 caused greatly reduced cell proliferation as well as significantly reducing the population of cells in S-phase. In a recent study, Lafarga et al. showed Tial1 to be essential for the G2/M checkpoint, accumulating in nuclear foci in late G_2_ and prophase in cells that experience replication stress. They found depletion of Tial1 to accelerate mitotic entry, leading to chromosomal instability in response to replication stress, in a manner that could be alleviated by the concomitant depletion of Cdc25B or inhibition of CDK1 (Lafarga *et al*, 2019). Although our results are not directly comparable with those of Lafarga et al., they are not incompatible, and the role of Tial1-containing subnuclear foci could be a valuable avenue of exploration in the elucidation of the mechanism by which Tial1 regulates DNA replication. These properties of Tial1 may also be reminiscent of a role in the formation of biomolecular nuclear condensates, which may regulate DNA replication as well as other nuclear processes (Sabari *et al*, 2020).

In conclusion, we have identified proteins that can bind the mouse OGRE and that could be potential players in regulating DNA replication origin activity, alone or in combination with each other. A common link between the factors we have characterised in this work, and consistent with a major enrichment for nucleic acid binding activity, is RNA binding, suggesting a possible implication of RNA in the regulation of DNA synthesis. In support of this possibility, growing evidence implicates different classes of RNA in regulating DNA replication (Kowalski & Krude, 2015; Aze *et al*, 2017; Marchese & Huarte, 2017; Zhang *et al*, 2023). Further studies will be important to elucidate the roles of these factors in this fundamental cellular process.

## 4. Materials and Methods

### Cell culture and extract preparation

NIH 3T3 immortalised mouse fibroblast and U2OS human osteosarcoma cell lines were cultured in Dulbecco’s Modified Eagle Medium (Gibco, Thermo Fisher), supplemented with 10% (v/v) foetal bovine serum (EUROBIO), 2 mM L-glutamine, 100 U/ml penicillin, 100 µg/ml streptomycin, in a 37°C incubator in an atmosphere containing 5% CO_2_.

### Nuclear extract preparation

Nuclear extracts of NIH 3T3 cells were prepared according to the method of Dignam (Dignam *et al*, 1983). Extracts were aliquoted, frozen in liquid nitrogen and stored at -80°C.

### DNA Affinity Purification

Isolation of proteins that associate with specific DNA sequences was performed based on a previously published method (Nordhoff *et al*, 1999). DNA oligonucleotides used for DAP-MS were synthesised by Eurofins Genomics. Biotin moieties, where present, were covalently attached to the 5′ end of one of the strands, with a triethyl glycol (TEG) spacer. The sequence of DAP probes was derived from the genome of mouse chromosome 11, surrounding the OGRE3 sequence.

The sequences of 80-mer probes were as follows:

OGRE3^WT^, Biotin-5′-CGAGGT TCCTAG GCGCCT AAAGAC CGAGTG GGGGCG GGGAGG GAAGGG GGTGCT GTGCGT GCGCGC GCGCGT GCCAAA GC-3′ and its complement 5′-GCTCCA AGGATC CGCGGA TTTCTG GCTCAC CCCCGC CCCTCC CTTCCC CCACGA CACGCA CGCGCG CGCGCA CGGTTT CG-3′.

OGRE3^MUT^, Biotin-5′-CGAGGT TCCTAG GCGCCT AAAGAC CGAGTG CCGAGC CGAGCT GGCGCC GCTGCT GTGCGT GCGCGC GCGCGT GCCAAA GC-3′ and its complement 5′-GCTTTG GCACGC GCGCGC GCACGC ACAGCA GCGGCG CCAGCT CGGCTC GGCACT CGGTCT TTAGGC GCCTAG GAACCT CG-3′.

The sequences of 120-mer probes were as follows:

OGRE3^WT^, Biotin-5′-GTTTCT TCGAAT TCCTGC TGCGAG GTTCCT AGGCGC CTAAAG ACCGAG TGGGGG CGGGGA GGGAAG GGGGTG CTGTGC GTGCGC GCGCGC GTGCCA AAGCTA GAAGGG AGGGAA CGCTTT-3′ and its complement 5′-AAAGCG TTCCCT CCCTTC TAGCTT TGGCAC GCGCGC GCGCAC GCACAG CACCCC CTTCCC TCCCCG CCCCCA CTCGGT CTTTAG GCGCCT AGGAAC CTCGCA GCAGGA ATTCGA AGAAAC-3′.

OGRE3^MUT^, Biotin-5′-GTTTCT TCGAAT TCCTGC TGCGAG GTTCCT AGGCGC CTAAAG ACCGAG TCGGCC AGGACC GGTGGC CGCCTG CTGTGC GTGCGC

GCGCGC GTGCCA AAGCTA GAAGGG AGGGAA CGCTTT-3′ and its complement 5′-AAGCG TTCCCT CCCTTC TAGCTT TGGCAC GCGCGC GCGCAC GCACAG CAGGCG GCCACC GGTCCT GGCCGA CTCGGT CTTTAG GCGCCT AGGAAC CTCGCA GCAGGA ATTCGA AGAAAC-3′.

To anneal the forward- and reverse-strand DNA oligonucleotides, each were mixed at 100 µM in annealing buffer (10 mM Tris-HCl pH 7.5, 50 mM NaCl, 1 mM EDTA). The mix was heated at 95°C for 5 mins in a thermocycler, then allowed to cool very slowly to room temperature, and stored at -20°C. Annealed oligos were bound to Streptavadin M-280 Dynabeads (Thermo Fisher, reference number 11205D), in annealing buffer, then washed with DAP buffer (10 mM HEPES-KOH, pH 7.9, 150 mM KCl, 1 mM MgCl_2_).

NIH 3T3 nuclear extracts (100 µl) were diluted with Dignam buffer A (10 mM HEPES-KOH, pH 7.9, 1.5 mM MgCl_2_, 10mM KCl, 0.5 mM DTT) to produce a final salt concentration of 150 mM. Extract was supplemented with ATP (1 mM), and LightShift™ poly(dI:dC) (ThermoFisher, 50 µg/ml) and sheared salmon sperm DNA (50 µg/ml) as competitors, added to the magnetic beads coupled to annealed DNA, and incubated with rotation for one hour at 4°C. Beads were recovered by magnetic capture, washed three times with 400 µl DAP buffer containing poly(dI:dC) sheared salmon sperm DNA (as above), plus 0.1% NP-40. Buffer was removed the beads, and proteins eluted by boiling for 5 mins in SDS-PAGE sample buffer (Laemmli, 1970).

To assess the yield and quality of the proteins isolated, 10% of the sample was separated on a NuPAGE™ 4-12% gradient Bis-Tris SDS-PAGE gel (Thermo Fisher) in MOPS running buffer (50 mM Tris, 50 mM MOPS, 1 mM EDTA, 0.1% (w/v) SDS). The gel was then stained using the Invitrogen™ SilverQuest™ Silver Staining Kit (Thermo Fisher).

### Sample preparation for MS

To prepare the remaining protein sample for MS analysis, the sample in SDS sample buffer was loaded onto a NuPAGE™ 4-12% gradient Bis-Tris SDS-PAGE gel, and migrated, in MOPS running buffer, approx. 0.5 cm into the gel. The gel was then stained with colloidal blue (Euromedex) for 1 hr. Gel slices that include the totality of the proteins loaded on a lane were excised. Each gel slice was further cut into pieces of approx 1 mm^3^, reduced by treatment with 10 mM DTT in 50 mM triethylammonium bicarbonate (TEABC), alkylated by treatment with 55 mM iodoacetamide in 50 mM TEABC, then digested by treatment with 6 µl of Trypsin Gold (Promega). Peptides were extracted and concentrated for MS analysis.

### LC-MS/MS analysis

LC-MS/MS was performed by the Functional Proteomics Platform (FPP), Montpellier. Peptides were separated on a nano-HPLC (Ultimate 3000, Dionex) with an inverse phase C18 Pepmap® column (Dionex, Thermo Fisher), following a gradient of 100% solution A (0.1% formic acid, 2% acetonitrile) to 40% solution B (0.1% formic acid in acetonitrile), 60% solution A, at a rate of 300 nl/min, over 80 mins. The HPLC was linked in-line with a nanoelectrospray source to an LTQ Orbitrap Elite mass spectrometer (Thermo Fisher). Spectra were captured using Xcalibur software, v2.2 (Thermo Fisher).

### MS data analysis, proteomic data processing

Spectral data were matched to entries from the UniProt mouse Reference proteome database (version 2016_04, UniProt Consortium) using the Andromeda algorithm (Cox *et al*, 2011) within the MaxQuant (v1.5.0.0) software suite (Cox & Mann, 2008), with carbamidomethyl cysteine as a fixed modification, and acetyl (N-terminal) and methionine oxidation as variable modifications. The false discovery rate (FDR) was set to 1% at the peptide and protein levels. For relative abundance measurement of proteins, label-free quantification (LFQ) values were determined using the MaxLFQ algorithm (Cox *et al*, 2014) within MaxQuant.

For the DAP-MS experiments involving the 80-bp probe, statistical analysis and graphing were performed using Perseus software (Tyanova *et al*, 2016).

For the DAP-MS experiments involving the 120-bp probe, a table of identified proteins and associated data was imported into Microsoft Excel for filtering and sorting. Protein entries for which peptides were identified for the no-DNA condition but not for the OGRE3^WT^ or OGRE3^MUT^ conditions were filtered out. Protein entries where the LFQ intensity was zero for all three conditions were filtered out. Protein entries for which the LFQ intensity for no-DNA condition was greater than OGRE3^WT^ and OGRE3^MUT^ were filtered out.

To identify non-specific proteins within the list, firstly lists of gene symbols corresponding to commonly-identified non-specific proteins were derived, from two publications (Trinkle-Mulcahy *et al*, 2008) (Hutchins *et al*, 2010). Matches between these non-specific genes and genes corresponding to proteins identified by MS were identified using Microsoft Excel, and these proteins were filtered from the list.

To identify proteins showing a greater abundance in the OGRE3^WT^ relative to the OGRE3^MUT^ condition, the LFQ intensities for OGRE3^MUT^ were subtracted from the LFQ intensities for OGRE3^WT^, and the proteins sorted by this difference. Also, for proteins for which the LFQ intensity for OGRE3^MUT^ was nonzero, the ratio of LFQ(OGRE3^WT^) to LFQ(OGRE3^MUT^) was calculated. Plots were generated in Microsoft Excel.

### Dataset exploration

#### Gene Ontology (GO) analysis

Gene symbols corresponding to the mouse proteins identified were used to identify the human orthologs of these proteins, for which greater functional annotation is available. Gene Ontology (GO) terms corresponding to the cellular compartment, molecular function and biological process were obtained from the UniProt database (https://www.uniprot.org/) (UniProt Consortium, 2023), and integrated into the main protein table.

#### Cell cycle functionality

A list of genes from the MitoCheck genome-wide RNAi screen whose knockdown in HeLa cells induced a cell division phenotype (Neumann *et al*, 2010) was obtained, and the gene symbols updated to the current ones by searching the Human Gene Nomenclature Committee (HGNC) database (https://www.genenames.org/) (Tweedie *et al*, 2021). This updated gene list was correlated with the genes corresponding to the proteins identified by DAP-MS.

#### Mouse knockout lethality

Each mouse gene symbol corresponding to proteins identified was used to interrogate the Mouse Genome Informatics (MGI) database (http://www.informatics.jax.org/) (Blake *et al*, 2021). Phenotypes of knockout mice were classified as shown in **Supp Table 1**.

#### Pathway analysis

UniProt accession codes corresponding to the proteins identified by DAP-MS were used to query the REACTOME database (https://reactome.org/) (Gillespie *et al*, 2022), and KEGG PATHWAY database (https://www.kegg.jp/kegg/pathway.html) (Kanehisa & Goto, 2000). Where a match was found to a known biochemical pathway, the corresponding graphic was exported.

### siRNAs

The siRNAs used to target the factors DHX36, PURA and TIAL1 were siGENOME SMARTpools, mixes of four distinct sequences (Horizon Discovery, PerkinElmer). Catalogue numbers were as follows: DHX36, M-013167-00; PURA, M-012136-01; TIAL, M-011405-01. The negative control siRNA was Silencer™ Select negative control N° 2 (cat# 4390846, Thermo Fisher).

### Antibodies

Primary antibodies used in this study were as follows:

| Target protein | Host organism | Supplier | Catalogue number |
| --- | --- | --- | --- |
| BrdU | Rat | Abcam | ab6326 |
| Dhx36 | Rabbit | Abcam | ab70269 |
| Pura | Rabbit | Abcam | ab125200 |
| Tial1 | Rabbit | Abcam | ab129499 |
| Tubulin | Mouse | Sigma-Aldrich | T6199 |

### Immunostaining and fluorescence microscopy

Cells cultured on coverslips were washed once in PBS, then fixed in 3% paraformaldehyde for 15 mins at room temperature. Cells were washed briefly with PBS, then incubated with 50 mM NH_4_Cl for 10 mins to saturate the aldehyde groups, then permeabilised with PBS + 0.2% Triton X-100 for 5 min. Cells were washed three times for 10 mins each in PBS + 0.5% BSA, then incubated for 1 hr with primary antibodies, diluted 1/100 in the same. After washing three times for 10 mins each in PBS + 0.5% BSA, coverslips were incubated in the secondary antibody: goat anti-Rabbit IgG, Alexa Fluor™ 488 conjugated (ThermoFisher, A11034), diluted 1/500 in PBS, 0.5% BSA), for 1 hr at room temperature. After washing three times for 10 mins each in PBS, the coverslips were left to dry at room temperature, then mounted in PBS + 75% glycerol. Images were captured using a Zeiss Axio Imager fluorescence microscope with ApoTome (structured illumination microscopy) attachment, and ZEN software (Carl Zeiss). Microscopy image acquisition was performed at the Montpellier Ressources Imagerie (MRI) platform in Montpellier, France (https://www.mri.cnrs.fr/en). Extremely low fluorescence signals were detected in control coverslips lacking the primary antibodies. Signal intensity values were determined by making measurements of ten nuclear or cytoplasmic regions for each condition, using ImageJ software (Schneider *et al*, 2012).

### Western blotting

Proteins boiled in sample buffer (Laemmli, 1970)) were separated on NuPAGE™ 4-12% gradient Bis-Tris SDS-PAGE gels, as described above. Proteins were electrophoretically transferred from the gel to a nitrocellulose membrane for probing using antibodies (Towbin *et al*, 1979). The membrane was Hybond-C nitrocellulose (GE Healthcare), and transfer was performed in 25 mM Tris, 200 mM glycine, 20% (v/v) ethanol, at 90 volts for 80 mins at 4°C. To assess the quality of protein transfer, the membrane was stained with 0.1% (w/v) Ponceau-S (Sigma-Aldrich), dissolved in 5% (v/v) acetic acid. The membrane was washed three times with Tris-buffered Saline Tween (TBST: 25 mM Tris-HCl, pH 7.6, 150 mM NaCl, 0.05% (v/v) Tween-20), then blocked with TBST containing 5% (w/v) non-fat milk (TBSTM), for 30 mins. Membranes were incubated in primary antibodies diluted in TBST plus 2% (w/v) BSA for 1 hr at 4°C. Membranes were washed three times with TBST, incubated in HRP-conjugated anti-IgG secondary antibodies (Cell Signaling Technology) in TBSTM, washed three times with TBST. Labelled proteins were revealed by chemiluminescence using Immobilon Classico, Crescendo (Merck Millipore), or SuperSignal™ West Femto (Thermo Fisher) reagents, and visualised using a ChemiDoc Imaging System (Bio-Rad).

Intensities of western blot bands were measured using ImageJ software (Schneider *et al*, 2012), with subsequent calculations performed using Microsoft Excel. For a given area including a band of interest, the mean greyscale value (0, black to 255, white) was first inverted then multiplied by the corresponding area to produce a total intensity. These values were corrected by subtracting the intensity of a region of identical area from a background region of the blot.

### Flow cytometry

Cell cycle profiles were determined by dual labelling of propidium iodide (PI) and a pulse of 5-bromo-2′-deoxyuridine (BrdU), followed by flow cytometry analysis (Sasaki *et al*, 1986). Cells were fixed, and nuclei prepared using a pepsin-digestion procedure, as described (Egger *et al*, 2022). Samples were analysed using a MACSQuant flow cytometer (Miltenyi Biotec) at the Montpellier Ressources Imagerie (MRI) platform. 10,000 nuclei were analyzed per sample. A forward scatter vs side scatter (FSC vs SSC) plot was used to gate out debris, and a PI Height vs PI Area plot was used to gate out doublets. Graphing and quantification of flow cytometry data were performed using FlowJo software v10 (Becton, Dickinson & Company).

## AUTHOR CONTRIBUTIONS

Conceptualisation, J.R.A.H. and M.M.; methodology, J.R.A.H., I.P., S.U., J.L.M., D.M. and M.M.; formal analysis, J.R.A.H., S.U.; investigation, J.R.A.H., I.P., S.U.; writing—original draft preparation, J.R.A.H.; writing—review and editing, J.R.A.H., D.M. and M.M.; visualisation, J.R.A.H., I.P. and S.U.; supervision, D.M. and M.M.; project administration, P.M., D.M. and M.M.; funding acquisition, J.R.A.H. and M.M. All authors have read and agreed to the published version of the manuscript.

## ACKNOWLEDGEMENTS

We thank Drs Christelle Cayrou, Philippe Coulombe, Tom Egger, Olivier Ganier and Sihem Zitouni for help, advice and useful discussions. Mass spectrometry experiments were carried out using the facilities of the Montpellier Proteomics Platform (PPM, BioCampus Montpellier). We are also grateful to Martial Seveno, Oana Vigy (Functional Proteomics Platform, Montpellier) for help with proteomic analysis, and Amelie Sarrazin and Marie-Pierre Blanchard (Montpellier Ressources Imagerie) for their assistance with flow cytometry and fluorescence microscopy, respectively.

---

**Supplementary Table 1.**
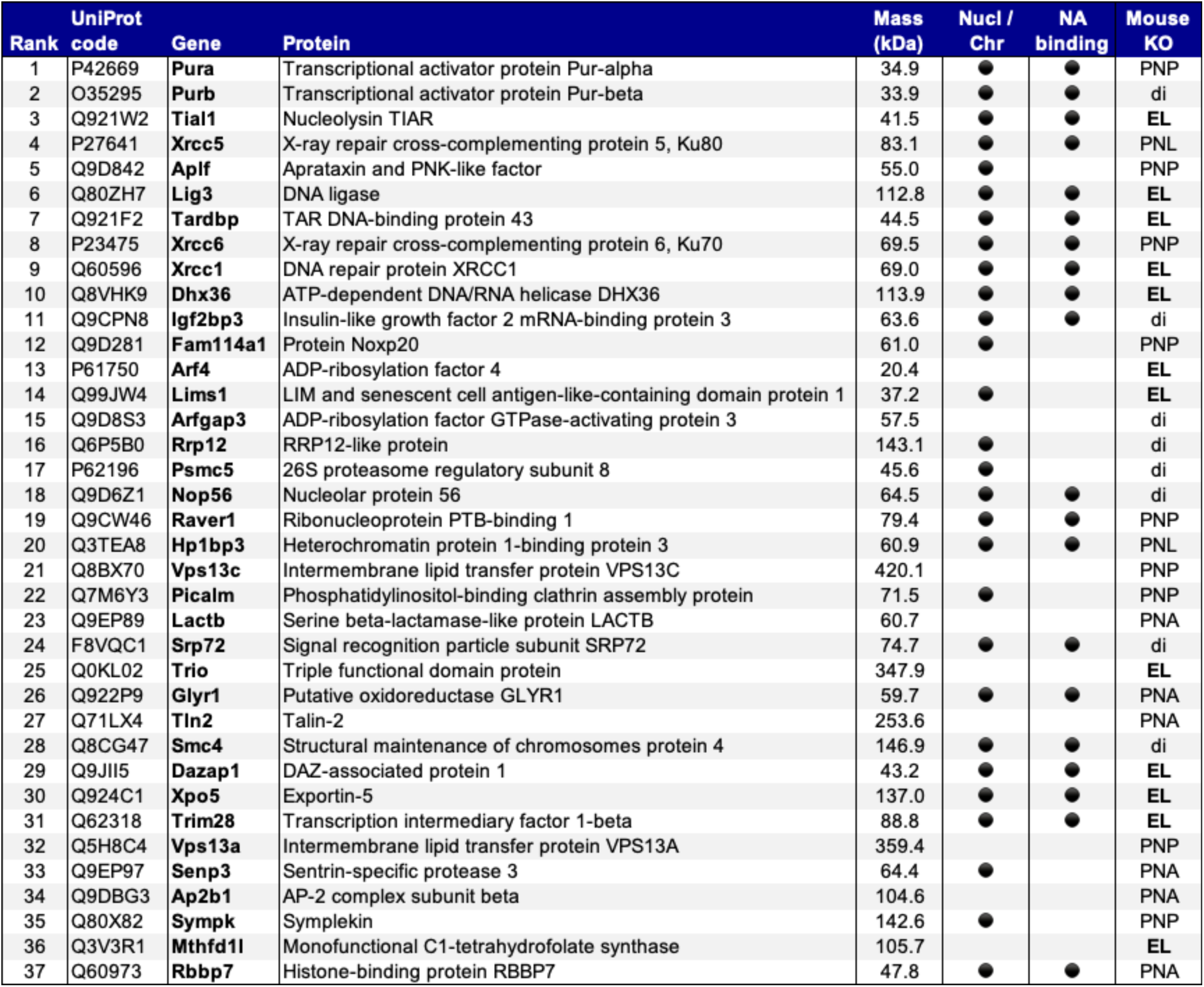
Proteins identified by DAP-MS as showing binding preference for OGRE3^WT^. These 37 proteins show a greater LFQ intensity for the 120-bp OGRE3^WT^ than for OGRE3^MUT^. Black circles indicate whether proteins are reported to localise to the nucleus or chromatin (Nucl/Chr), and contain a nucleic acid binding domain (NA binding). Also shown are reported consequences of mouse knockout (Mouse KO): EL, embryonic lethal; PNA, postnatal abnormalities; PNL, postnatal lethal; PNP, postnatal pathology; di, data insufficient.

**Supplementary Figure 1.**
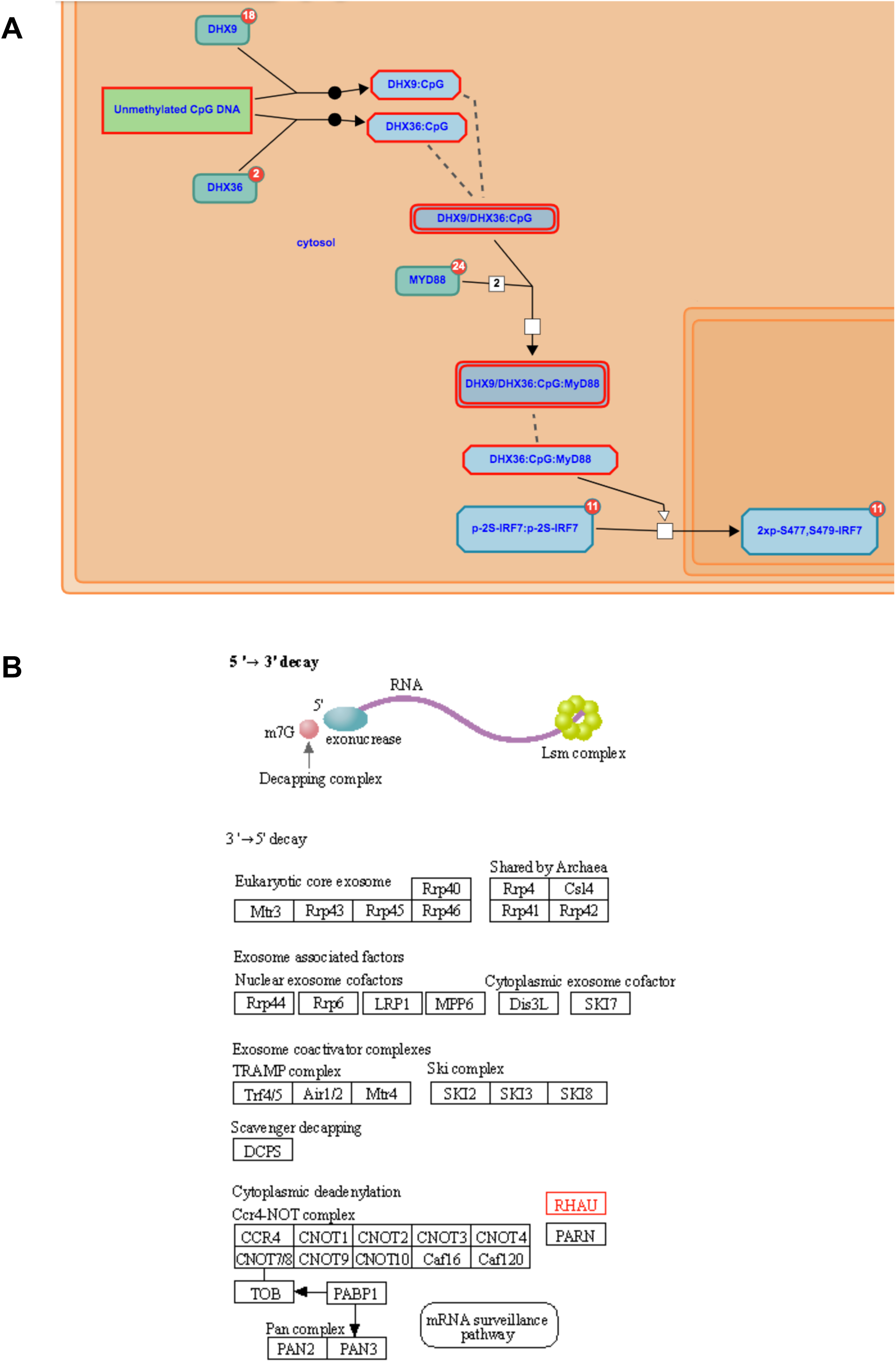
– Pathway analysis of Dhx36. (A) Role of Dhx36 in a pathway that acts as a cytosolic sensor of pathogen-associated DNA as part of the innate immune system (from REACTOME). **(B)** Role of Dhx36 (RHAU) in RNA degradation (KEGG Pathway, map03018).

**Supplementary Figure 2.**
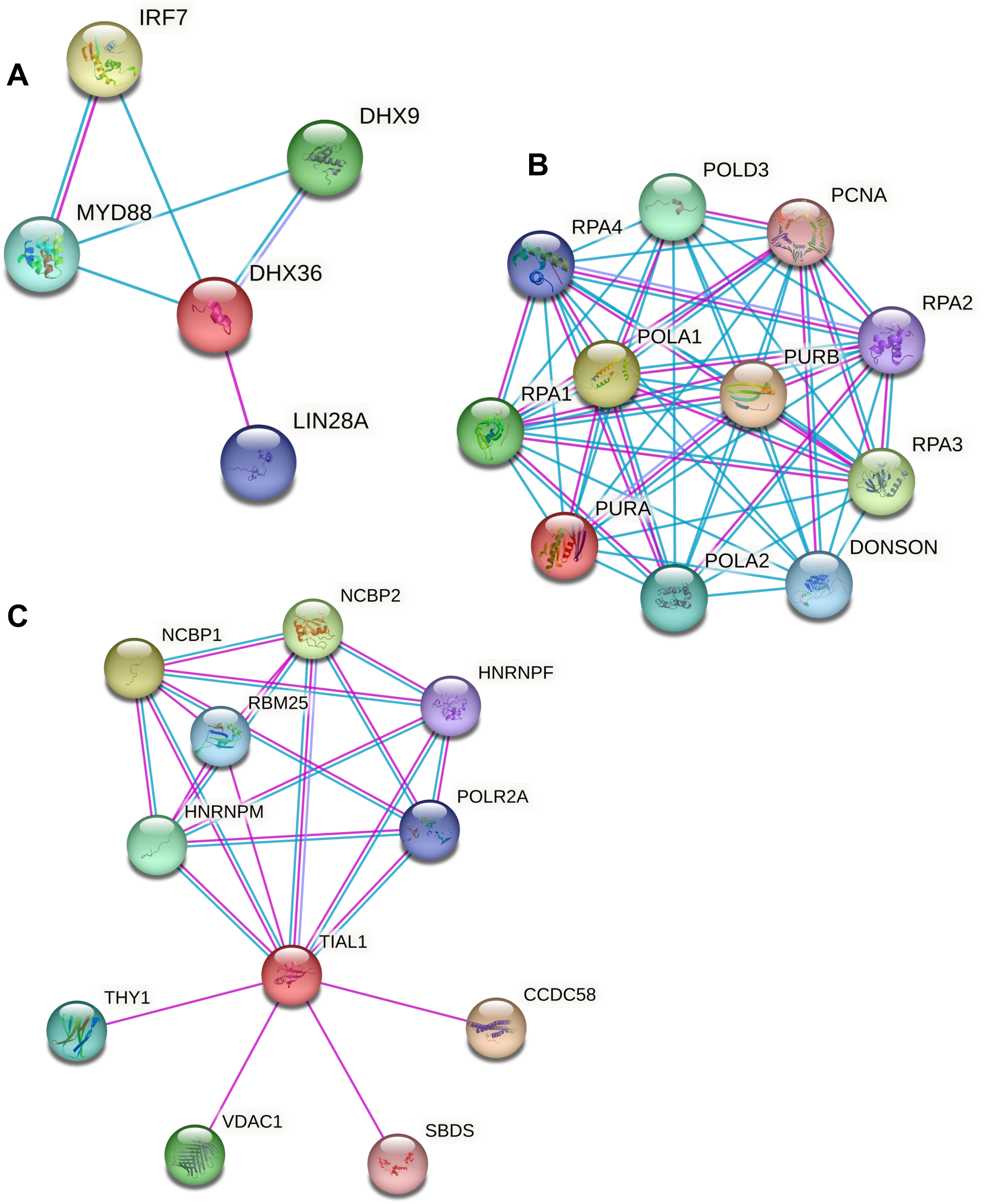
– STRING interaction graphs of the three candidate factors. Interaction graphs centred on **(A)** Dhx36, **(B)** Pura, and **(C)** Tial1 are shown. Coloured nodes indicate the respective proteins, labelled according to official gene symbols. Magenta edges represent interactions derived from “experiments”, cyan edges represent interactions from “databases”.

**Supplementary Figure 3.**
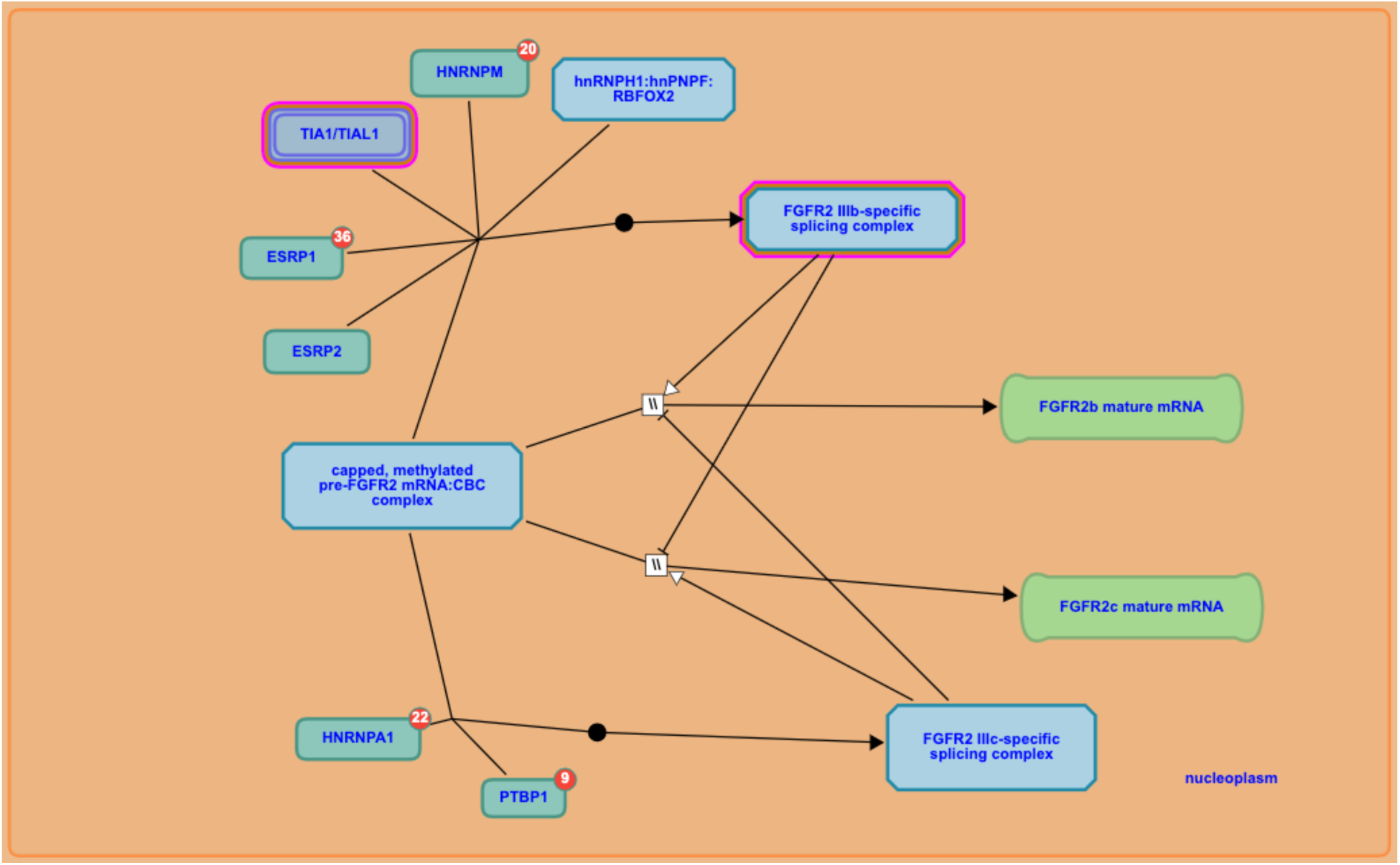
– Pathway analysis of Tial1. Tial1, with Tia1, ESRP1 and 2, RBFOX2 and HNRNPM contribute to the formation of the FGFR2 IIIb splice variant, characteristic of epithelial cells (from REACTOME).

